# Have coral snake mimics diversified more than non-mimics?

**DOI:** 10.1101/042440

**Authors:** Daniel S. Caetano, Laura R. V. Alencar, Paulo Passos, Felipe G. Grazziotin, Hussam Zaher, Marcio Martins

## Abstract

Dipsadidae is the most diversified family of snakes, composed of species showing an impressive variety of color patterns. Some species are cryptic whereas others have contrasting patterns comprised by bright colors alternated with darker shades, including particular combinations of vivid colors characteristic of coral snakes (Elapidae). Species with such patterns are thought to be mimics of coral snakes based on their color pattern similarity, predator avoidance of such patterns in field experiments, and the geographical concordance between models and mimics. Here we test whether color patterns associated with coral snake mimicry and contrasting color patterns in general influenced the diversification dynamics of the group. We compile the largest database of color patterns among reptiles to date, with color descriptions for the majority (594 species) of dipsadids. We used trait-dependent diversification models along with extensive simulations to deal with the recently described statistical bias associated with such methods. Despite the apparent survival advantage associated with coral snake mimicry, we show that there is no detectable influence of color types in the dynamics of diversification in Dipsadidae. We discuss insights into the function of color patterns and argue that non-mimic contrasting patterns might serve as pre-adaptations to mimicry of coral snakes.

**Data archival location:** BEAST XML file and BiSSE MCMC results: http://dx.doi.org/10.6084/m9.figshare.831493

R code for analyses and simulations: https://github.com/Caetanods/Dipsadidae_color_evolution

---

Colors play an important role in avoiding predation. Patterns similar to the background environment make prey difficult for the predators to detect and recognize (Merilaita and Lind 2005, Stevens and Merilaita 2009). On the other hand, bright and contrasting colors displayed by unpalatable, toxic or venomous animals (i. e., aposematic patterns) serve as warning signals that are often avoided by visually oriented predators (Wallace 1867, Mappes et al. 2005, Speed and Ruxton 2005). However, such conspicuous colors can also be displayed by mimics, which gain protection by deceiving predators that avoid their false warning signals. Strong evidence from field experiments shows that mimicry of warning signals decreases predation pressure when compared to cryptic color patterns (Jeffords et al. 1979, Brodie 1993, Brodie and Janzen 1995, Pfennig et al. 2001, Pinheiro 2011, Pfennig et al. 2015). Such reduction in predation pressure may also have positive impacts on habitat use by aposematic lineages and their mimics. Cryptic animals are to some degree restricted to backgrounds which their color patterns match and may only be active at certain times because movement is often antithetic to good crypsis (Speed et al. 2010, Stevens and Ruxton 2012). In contrast, such restrictions may be weaker in aposematic or mimic lineages, which could promote more opportunities to exploit habitat resources (Speed et al. 2010).

In contrast with aposematism, the survival advantage of Batesian mimicry is dependent on the relationship between the model and the mimetic organism because predators need to associate the unpalatability or hazard of the model with the warning signals of the deceiver. Once this association is broken, a mimicry breakdown occurs and the mimic phenotype might become maladaptive since warning signals can make individuals more conspicuous to predators (Mallet and Joron 1999, Pfennig et al. 2001, Pfennig et al. 2015). Mimicry breakdown can be caused by allopatry between mimic and model populations as a result of population expansion of the mimic or local extinction of the model (Pfennig and Mullen, 2010). Allopatric mimics are conspicuous to naïve predators that might not avoid their deceptive warning signals and this may result in higher predation rates and eventual extinction of the mimic population (Pfennig et al. 2015). On the other hand, population expansion or migration of mimics can create opportunities for local adaptation to novel aposematic models. This process could result in selection against intermediate hybrids followed by decreased gene flow among populations and eventually promote reproductive isolation (Mallet and Joron 1999, Pfennig et al. 2015). Over longer time scales such processes might have a positive effect on rates of diversification of mimetic lineages. Previous studies show that aposematic lineages are more species-rich than cryptic ones (Santos et al. 2003, Przeczek et al. 2008), suggesting that the evolution of the aposematic condition may even represent a key innovation (Speed et al., 2010). This key innovation hypothesis could be extended to mimicry; however, the potential effects of mimicry evolution on lineage diversification have yet to be investigated.

Among snakes, groups of relatively harmless or mildly venomous species showing color patterns similar to those of venomous coral snakes (Elapidae) have instigated a long debate on whether such patterns are mimetic (see a comprehensive review in Pough 1988). Savage and Slowinski (1992) compiled a remarkable list of coloration descriptions and designed a system of categories to facilitate recognition of coral snake mimics and association with their supposed models. Early reports also relied primarily on the similarity of color patterns between mimics and models to argue in favor of mimicry relationships (Dunn 1954, Hecht and Marien 1956, also see Greene and McDiarmid 1981). Additional evidence came from parallel geographic variation of coral snakes and their putative mimics (e. g., Hecht and Marien 1956, Zweifel 1960, Greene and McDiarmid 1981, Marques and Puorto 1991) and from field studies using replicas of coral snakes and other similar color patterns (Smith 1975, Brodie 1993, Brodie and Janzen 1995, Hinman et al. 1997, Pfennig et al. 2001, Buasso et al. 2006).

Some authors pointed to the possibility that contrasting colors, including the stereotypical banded pattern observed in almost all coral snakes, could serve a disruptive function (Gadow 1908, Thayer 1918, Dunn 1954, and Brattstrom 1955). Those reports suggested that the alternate pattern of bands could blend to the background environment and break the outline of the snake body, making recognition by visually oriented predators difficult. Recently, Titcomb and colleagues (2014) showed that the contrasting ringed pattern of coral snake mimics can create an illusory effect when the individuals are moving fast. The effect, called flicker-fusion, can give advantage to snakes against avian predators independent of mimicry. Despite its protective effect, the plausible disruptive function of the contrasting bands do not invalidate the existence of a mimicry complex between elapids and snakes from other families, since the same color pattern can perform both functions (Titcomb et al., 2014).

The family Dipsadidae (Zaher et al. 2009, sensu Grazziotin et al. 2012) is the most diverse among snakes, with ca. 700 species occurring from Central to South America (Grazziotin et al. 2012, Uetz and Hosek 2014), and is characterized by an impressive variety of color patterns (see Martins and Oliveira 1998 for some examples). Some dipsadids have color patterns similar to those of coral snakes, and have long been suggested as cases of mimicry of New World coral snakes of the genus *Micrurus* and *Leptomicrurus* (family Elapidae; Wallace 1867, Greene and McDiarmid 1981, Sazima and Abe 1991, Savage and Slowinski 1992, Martins and Oliveira 1993, Pough 1988, Almeida et al. 2014). The contrasting coloration found in dipsadid snakes always includes bright colors but is not restricted to ringed patterns. In general, species can vary from the coral snake pattern of black, red and yellow rings or bands to a less colorful homogeneous red body with a single black or cream band on the neck (nuchal collar). Besides contrasting color patterns, the family also shows a diverse array of cryptic color patterns, characterized by blotches and shades of brown, gray, or green. Included in the latter are species whose dorsum is cryptic and whose venter has a plain bright color and even a coral snake pattern. Mimetic and cryptic patterns can be found both within and among genera and make dipsadid snakes an ideal study system to investigate the possible effects of such distinct color types on macroevolutionary patterns.

Herein we test whether distinct color patterns have an influence on the diversification of the family Dipsadidae. We investigate whether color patterns similar to coral snakes (and contrasting color patterns in general) show diverging macroevolutionary patterns when compared to non-mimic and cryptic lineages, respectively. We compile and make available a database with color pattern descriptions for the vast majority of species in the group. We show that there is no detectable influence of supposedly mimic or contrasting color patterns in the dynamics of diversification and argue that non-mimic contrasting color patterns may be pre-adaptations to mimicry of coral snakes.

## Methods

### Phylogenetic reconstruction

We used sequence data for Dipsadidae and outgroup species available in GenBank (Benson et al. 2014) and previously analyzed by Grazziotin and colleagues (2012, see accession numbers in their Appendix S1). We aligned sequences using MAFFT (Katoh et al. 2005) under the G-INS-i strategy and selected models of molecular evolution for each of the eight gene sequences using a decision theory framework in DT-ModSel (Minin et al. 2003). We concatenated the alignments and set four partitions; one partition for each nuclear gene (bdnf, c-mos, and rag2) and a single partition with the mitochondrial genes (12S, 16S, cytb, nd2, and nd4). We used phyutility (Smith and Dunn 2008) to trim down all sites with 75% or more missing data and inferred a Maximum Likelihood (ML) tree using GARLI 2.0 (Zwickl 2011). We used the resulting ML phylogeny as the starting tree for three independent searches in BEAST 1.8 (Drummond et al. 2012) for 270 million generations with a thinning interval of 1500 generations each. Since there are sequences available for only few species of each genera we set an incomplete sampling birth-death tree prior (Stadler 2009) and an uncorrelated relaxed clock model to estimate relative branching times. We checked each run for convergence using Tracer 1.6 (Drummond et al. 2012) and excluded 50% of the posterior chain as burnin. We then combined the posterior set of trees from the three BEAST searches and randomly sampled 100 trees to account for phylogenetic uncertainty in all subsequent analyses. The 100 sampled trees and the BEAST xml file comprising the data matrix, selected models of molecular evolution, starting tree and prior parameters is available in FigShare (http://dx.doi.org/10.6084/m9.figshare.831493). We also deposited the configuration and log files for GARLI 2.0. Figure S1 shows the resulting maximum clade credibility (MCC) tree and respective posterior probability support values.

### Color patterns

To understand the evolution of colors and its effect on diversification we compiled the most complete database of coloration patterns for dipsadid snakes. We searched several information sources such as comprehensive taxonomic reviews (e.g., Downs 1967), published articles and books containing photographs of identified individuals (e.g., Savage 2002), trusted on-line photo repositories (e.g., CalPhotos - http://calphotos.berkeley.edu/ and Reptile Database – http://www.reptile-database.org/), photographs of live specimens, and examination of individuals preserved in scientific collections (e.g., type specimens). We excluded invalid taxa or names presenting nomenclatural problems that are still appearing in the literature or online databases. We avoided subspecific ranks for coding the currently recognized taxa (with the exception of four subspecies of *Alsophis antillensis*) because terminals in available phylogenies correspond to species only and less than 10% of the members of the family Dipsadidae present valid subspecies to date.

While color diversity makes the family Dipsadidae interesting for studies focusing on the evolution of color patterns such as ours, this is also the most challenging characteristic of the system. Since it is not possible to consider all diversity of color patterns for comparative analyses, we used broad categories that are directly related to the hypotheses tested. In a first test, we used the categories *coral-mimic* and *non-mimic* (see Figure 1 and database available in http://dx.doi.org/10.6084/m9.figshare.831493). We call *coral-mimics* species that resemble the pattern of any New World coral snake species (see Roze 1996, Campbell and Lamar 2004). Species included in this category can show the coral-mimic pattern throughout the dorsum (e.g., *Simophis rhinostoma*) or restricted to the anterior portion of the body (e.g., *Pseudoboa coronata*). It is impossible to elect species-specific mimicry hypotheses for all 121 species included in the coral-mimic category given our current knowledge of the ecology and geographical distribution of the group. Therefore, we assigned species based on their color pattern similarity with putative models. On the other hand, all species not defined as potential mimics of coral snakes, independent of whether their color pattern was better described as contrasting or cryptic, were included in the category of *non-mimics*. As a result, the *non-mimic* category comprise species with cryptic color patterns and others with bright coloration but not resembling any known lineage of New World coral snake.

**Figure 1.**
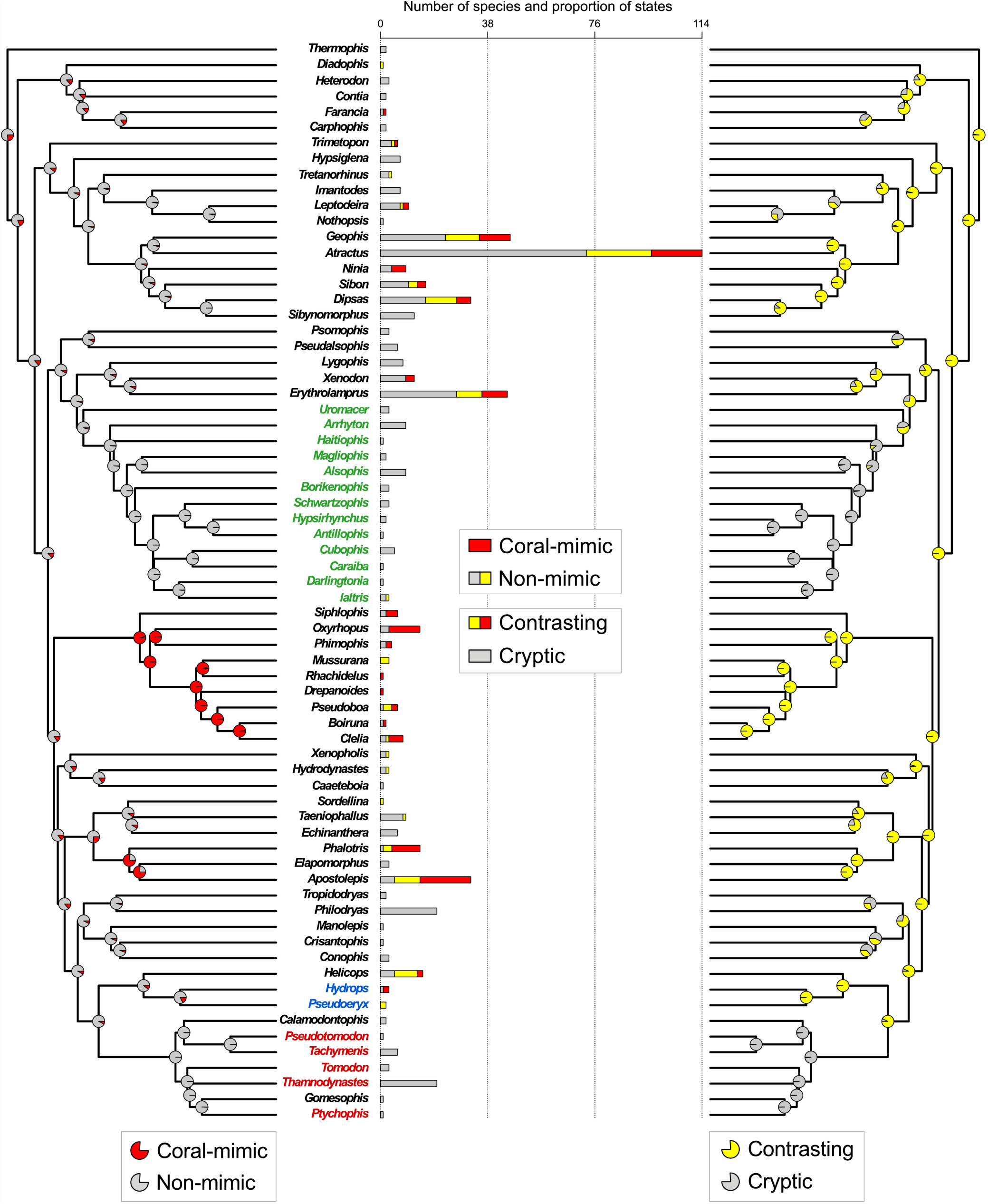
Maximum clade credibility (MCC) tree of the family Dipsadidae showing the number of species assigned to each color category (center) and the Maximum Likelihood (ML) ancestral state estimate for the color categories: *coral-mimic* versus *non-mimic* (left) and *contrasting* versus *cryptic* (right). Each color of the stacked bar chart (center) correspond to characteristics of the color patterns; species that do not show contrasting color patterns (i.e., cryptic coloration) are shown in gray, species with contrasting color patterns but that are not supposed mimics of coral snakes are shown in yellow, and species that show contrasting color patterns and are considered mimics of coral snakes are shown in red. The legend in the center of the plate applies different combinations of gray, yellow, and red to show the number of species assigned to the color patterns: *coral-mimic* (red) versus *non-mimic* (gray and yellow) and *contrasting* (yellow and red) versus *cryptic* (gray). The phylogeny on the left shows the ML ancestral estimate for *coral-mimic* in red and *non-mimic* in gray. The phylogeny on the right shows the ML ancestral estimate for *contrasting* in yellow and *cryptic* in gray. Both reconstructions were made under the trait-independent BiSSE model. The colored genera correspond to those associated with a tendency for higher speciation rates as a result of the MEDUSA analyses (compare with Figure S3). The tribe Alsophiini is highlighted in green, the two genera belonging to the tribe Hydropsini are marked in blue, and the taxa in red belong to the Tachymenini. Support for the nodes of the MCC tree are provided in Figure S1.

Our definition of *non-mimic* species includes both species that show cryptic color patterns such as hues and blotches of brown or green and bright colored patterns, which are likely to be more conspicuous to a visually oriented predator. One might argue that such definition is too inclusive, and the same category comprises different defensive strategies. In this light, we performed separate analyses using species that show brightly colored patterns in general, independent of whether the color pattern was similar to those of coral snakes. We defined such lineages as *contrasting*. Given our definitions, the category *coral-mimic* is a subset of the *contrasting* category; every *coral-mimic* lineage is among the species defined as *contrasting*, but the reverse is not true. We included species not classified as *contrasting* into a single category comprising *cryptic* patterns. Cryptic coloration has been defined as any pattern resembling a random sample of the habitat background (Endler 1986). However, since we do not have accurate habitat descriptions for most of species, we defined as *cryptic* all color patterns lacking contrasting colors (with exception of arboreal snakes, see below). Examples of such patterns are blotches with hues of brown, reddish brown, gray, and other combinations of dark colors. We also considered as *cryptic* species whose dorsum is homogeneously green, since individuals of those species are usually found among leaves of trees and bushes (e.g., *Uromacer*). Some Pseudoboini snakes (sensu Zaher et al. 2009) show ontogenetic changes in color pattern in which juveniles are brightly colored but become cryptic when adults (Martins and Oliveira 1998). We included those species into the *contrasting* category since juveniles correspond to the life stage most threatened by predation (Bonnet et al. 1999) and thus their defensive tactics are fundamental for individuals to reach sexual maturity. Some species show cryptic color patterns in the dorsum but have contrasting patterns restricted to the venter. The distinct patterns in the dorsum and venter are usually associated with a threatening display in which individuals twist the body and expose the bright colors when disturbed (Martins and Oliveira 1993, Sawaya et al. 2008, Tozetti et al. 2009). Hence, we classified those cases into the *contrasting* category instead of following their dorsal coloration. We performed additional analyses in which those species were classified as cryptic, but found no appreciable difference in results.

Some species are known to show color polymorphism. In such cases populations may show cryptic patterns occasionally associated to thermoregulation (Tanaka 2005, but see Lorioux et al. 2008). Alternatively, contrasting colors in polymorphic populations can be due to increasing sexual dichromatism in the course of the reproductive season (Forsman 1995, Lindell and Forsman 1996), related to non-selective processes, such as migration and dispersal (King and Lawson, 1995) or genetic drift in local (Brakefield, 1990) or island populations (Bittner and King, 2003). Independent of the potential sources of the polymorphic color patterns we assigned species to the *cryptic* category every time a cryptic morph was described among the color types.

### Comparative analyses

We sampled 100 trees from the posterior distribution of the BEAST analysis, rescaled all trees to a total depth of 1, and retained only one randomly selected species of each genus while pruning the rest. When the original tree had paraphyletic genera we selected the most inclusive monophyletic clade representing each group and kept a single species to represent each of those genera. We used the terminally unresolved trees (FitzJohn et al. 2009), the number of species of each genus and their color patterns to test hypotheses of trait-dependent diversification using the Binary State Speciation and Extinction model (BiSSE - Maddison et al. 2007, FitzJohn et al. 2009, FitzJohn 2012). We repeated all BiSSE analyses based on the two color categorizations: *coral-mimic* versus *non-mimic* and *contrasting* versus *cryptic*. We used Markov chain Monte Carlo (MCMC) to estimate the posterior distribution of the parameters for the unconstrained BiSSE model (six free parameters), in which speciation (**λ**) and extinction (**μ**) are estimated dependent on the color pattern of the lineages, and the constrained model, in which **λ** and **μ** are not related to color types (four free parameters). We used an exponential prior distribution with rate parameter equal to 0.3 for all BiSSE parameters and a starting point equal to the maximum likelihood estimate (MLE). We ran 10,000 generations of MCMC (chain length was based on preliminary analyses), discarded 50% of generations as burnin and checked convergence using the ‘coda’ package (Plummer et al. 2006). To test whether the full model explained the data better than the trait-independent model we performed model selection using the Bayesian Deviance Information Criteria (DIC – Gelman et al. 2014). We performed all analyses with both the *coral-mimic* versus *non-mimic* and the *contrasting* versus *cryptic* categories.

In addition to the BiSSE model, which estimates homogeneous diversification rates across the tree, we used Medusa (Alfaro et al. 2009) to test whether rates of diversification change independent of color types. This model estimates rates of diversification and jump locations in a given phylogeny using a stepwise AIC approach to sequentially fit models of diversification with increased number of jumps in rates of speciation and extinction. In order to incorporate the effect of uncertainty in topology and branch lengths we applied Medusa across all 100 sampled trees. However, there is no simple solution to summarize those results, since distinct tree topologies have unique sets of nodes. Thus, we chose to focus only in the rate estimates for the external branches (i.e., the genera). This approach allows us to compare groups of genera showing consistently higher or lower rates of diversification with respect to the background rate, ignoring information about the specific position of each jump.

Recently, Rabosky and Goldberg (2015) showed that the BiSSE model is likely to produce significant results even when there is no true relationship between traits and shifts in diversification rates. We performed simulations to correct our results for this deviation. Since the BiSSE model influences the dynamics of diversification and makes it difficult to simulate trait data under a fixed tree, each simulation generated both trait data and phylogeny. First, we did posterior predictive checks for model adequacy to test whether data simulated under the trait-dependent and trait-independent BiSSE models are similar to the observed data. For each model, we simulated traits using the ‘tree.bisse’ function in the package ‘diversitree’ (FitzJohn 2012) with parameters drawn from the joint posterior distribution resulting from the BiSSE analysis and constraining simulations to have a tree depth equal to 1 (identical to the empirical trees). Finally, we compared the number of species and the relative frequencies of each trait generated by the simulations with the empirical dataset. If models are adequate, simulated phylogenies should produce both diversity and frequency of states similar to the observed data.

In order to check if the support for the chosen model is not a statistical artifact, we created 100 new datasets by drawing from a binomial distribution with probabilities equal to the frequency of each trait in the observed data. We sampled 10 trees from the posterior distribution that resulted from the BEAST analysis and performed an MLE of the BiSSE model (trait-independent and full model) using each of the simulated datasets. We computed the likelihood ratio test for each BiSSE estimate and produced a distribution of p values for each of the 10 sampled trees. This distribution represents the expected p values for the likelihood ratio test when traits have no effect in diversification. Following, we tested whether p values obtained in the model test using the observed data and the same trees are significantly smaller than the distribution under the null model (i.e., data generated from a binomial distribution).

To perform ancestral state estimates, we calculated the grand mean of the posterior distribution of transition rates (**q_01_** and **q_10_**) across all 100 sampled trees under the accepted model after performing the BiSSE simulations. Then we used those values to infer the MLE of the marginal ancestral states reconstruction on the maximum clade credibility (MCC) tree. We performed all comparative analyses in R (R Core Team 2015). Scripts to replicate analyses, simulations, and figures are available in https://github.com/Caetanods/Dipsadidae_color_evolution.

## Results

We compiled the largest report of coloration descriptions for reptiles comprising data for 594 species of dipsadid snakes and covering over 80% of the known diversity of the group. We were able to get detailed color descriptions for most species, but for some the information available was incomplete or limited (i.e., taxa known from a single specimen). Although those cases were not suitable for definition as *coral-mimic* or *non-mimic* lineages (i.e., state unknown or ‘NA’), we managed to classify those as either *contrasting* or *cryptic* for all but a few exceptions. Among all data sources, museum specimens are the most difficult to categorize since colors fade after preservation and only light and dark hues remain. Bright colors such as yellow, orange, pink, and red (derived from carotenoid pigments) fade completely over time turning into cream on preservative fluid. In contrast, dark pigmentation is preserved and sometimes turns into shades of black or dark brown. As a result, we assigned museum specimens with alternate bright and dark bands (or with a distinct nuchal collar) as *contrasting* and homogeneous light or dark patterns as *cryptic*. We provide the species list and their color patterns under both categorization schemes in the online data repository (https://github.com/Caetanods/Dipsadidae_color_evolution).

We used the BiSSE model to investigate whether color patterns have an effect on the diversification rates of the snake family Dipsadidae. We made analyses based on two different categorizations of the same dataset: *coral-mimics* versus *non-mimics*, which tested for differences in macroevolutionary patterns between lineages with and without color patterns similar to New World coral snakes, and *contrasting* versus *cryptic* color patterns, which investigated changes in diversification associated with the presence or absence of bright coloration. When we compared *coral-mimic* to *non-mimic* patterns, the BiSSE analysis showed that *coral-mimic* lineages have net diversification rates (**λ_1_** - **μ_1_**) in average two times higher than *non-mimics* (Figure 2). When diversification rates are constrained to be independent of color types we recovered intermediate net diversification values relative to the trait-dependent model. These results are similar to the analyses based on *contrasting* versus *cryptic* color patterns (Figure 2). Both the trait-independent and trait-dependent diversification models estimated strongly asymmetrical transition rates with changes from the *coral-mimic* to the *non-mimic* state (**q_10_**) in average two to three times more frequent than the reverse (**q_01_**). Median transition rates are qualitatively comparable independent of color categorization (Figure 3). On the other hand, the posterior distributions of transition rates for *coral-mimics* versus *non-mimics* are wider than *contrasting* versus *cryptic*, showing more uncertainty in parameter estimates. Results were consistent across all sampled trees and in all cases the state-dependent diversification model was the one preferred by the DIC model selection criteria (Figure S2).

**Figure 2.**
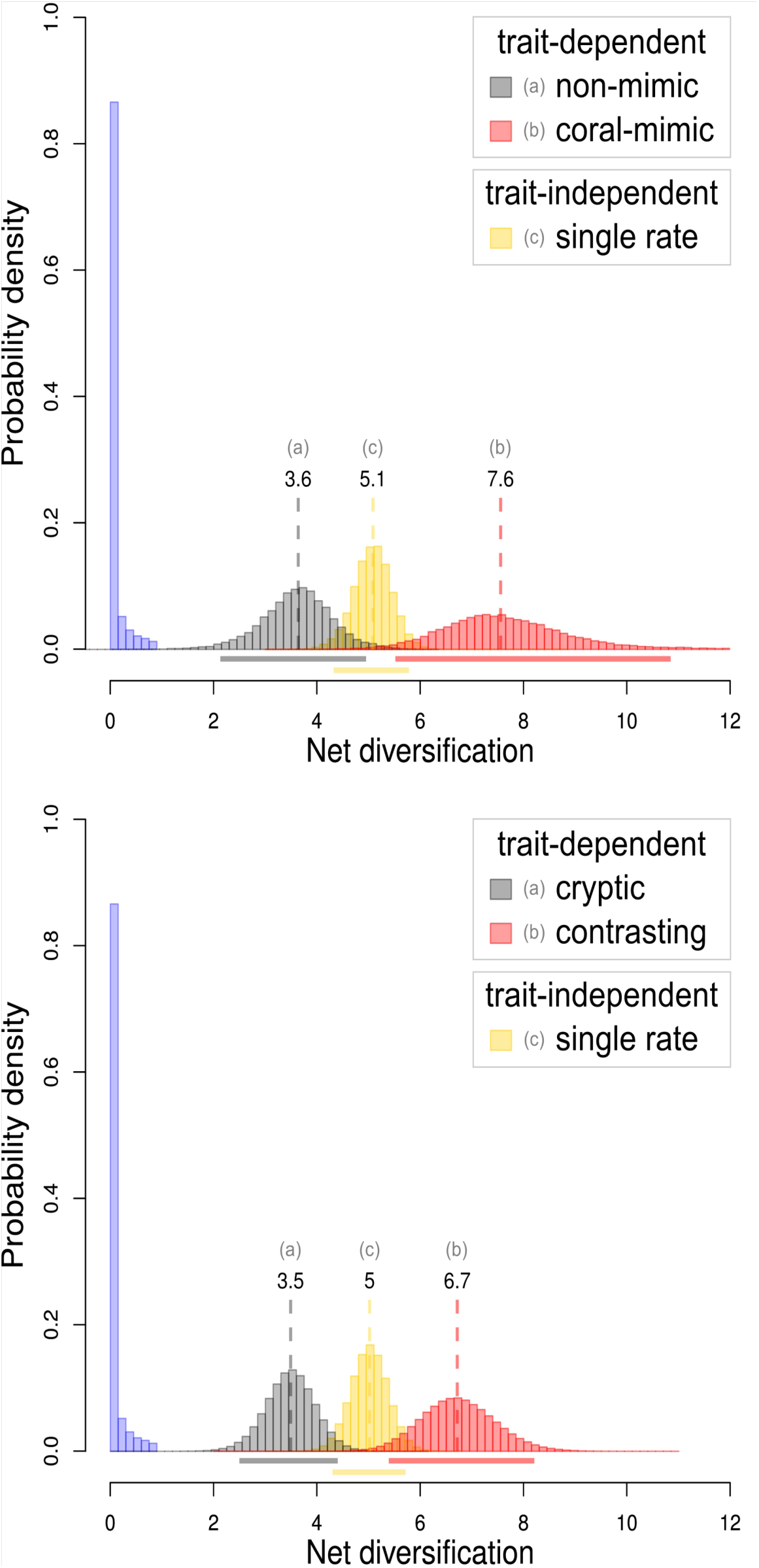
Posterior distributions of net diversification rates under the trait-dependent and trait-independent BiSSE models. Estimates are the joint posterior distribution from 100 MCMC BiSSE runs with randomly sampled trees. Prior distributions for the MCMC searches are shown in blue (unmarked distributions), the horizontal lines below each posterior distribution represent the 95% confidence interval, and the vertical hashed lines show median values. Top: Trait-dependent diversification rates of the *non-mimic* color pattern as gray and the *coral-mimic* color pattern as red. Bottom: Trait-dependent model with *cryptic* color pattern in gray and *contrasting* pattern in red. Both charts show the single diversification rate of the trait-independent model in yellow. Each posterior distribution is also identified by the letters (a), (b), and (c).

**Figure 3.**
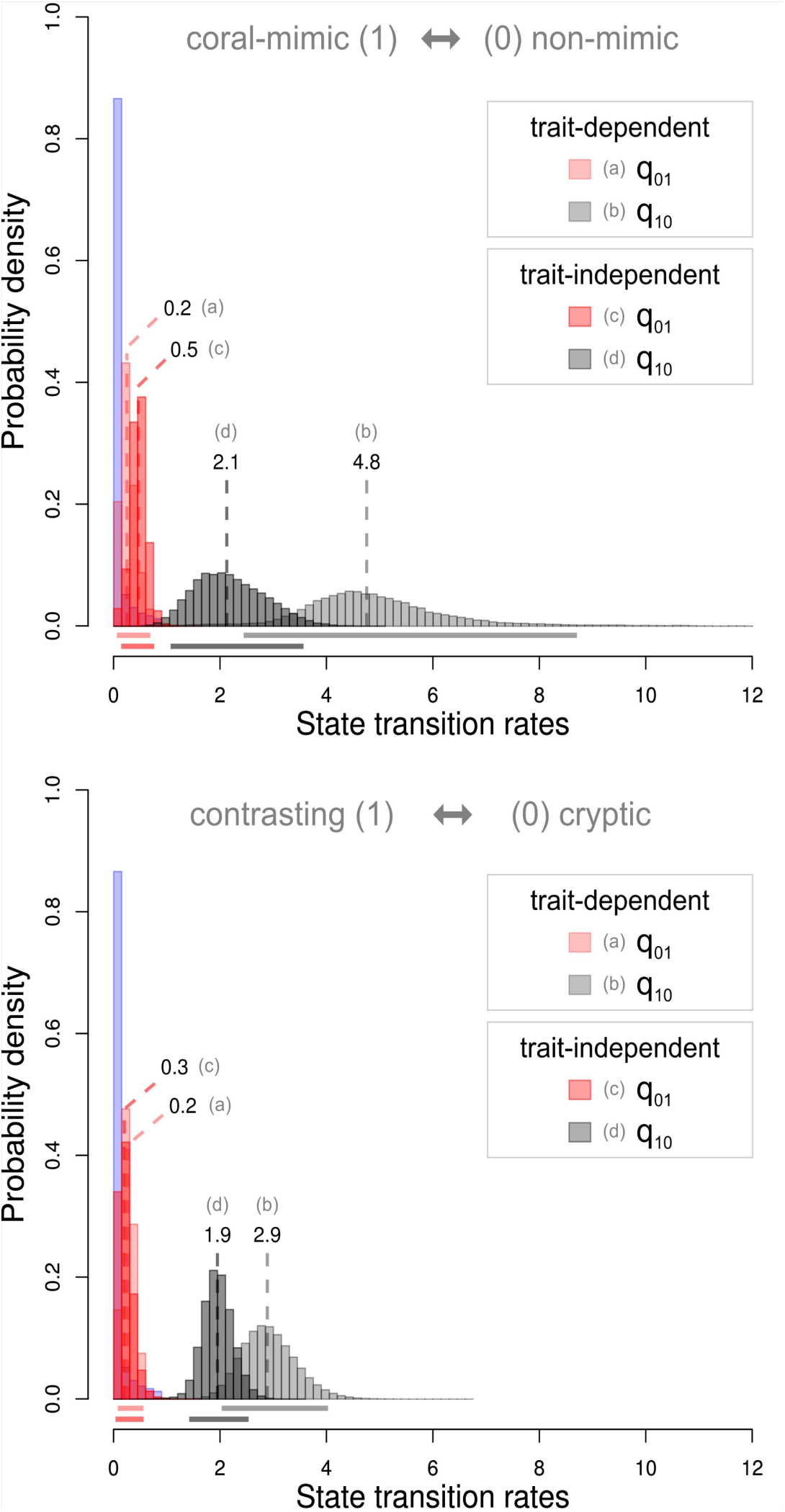
Posterior distributions of transition rates under the trait-dependent and trait-independent BiSSE models. Estimates are the joint posterior distribution from 100 MCMC BiSSE runs with randomly sampled trees. Prior distributions for the MCMC searches are shown in blue (unmarked distributions), the horizontal lines below each posterior distribution represent the 95% confidence interval, and the vertical hashed lines show median values. Top: Transition rates from the *non-mimic* to the *coral-mimic* (q01 - red) color pattern and back (q10 - gray). Bottom: Transition rates from the *cryptic* to the *contrasting* (q01 - red) color pattern and back (q10 - gray). Each chart shows results from the trait-dependent and trait-independent BiSSE models identified with different shades of the correspondent transition color and the reference letters (a), (b), (c), and (d).

Results from the Medusa analyses, which estimate shifts in diversification rates independent of color types, indicate that the preferred model across all 100 trees was the Yule diversification model (Yule 1924), which has a single birth parameter in the absence of extinction. The estimated median rate is comparable across almost all genera, but some clades have a tendency for higher rates of diversification when compared to background rates (Figure S3). While most of the 100 sampled trees were estimated to have homogeneous rates of diversification, a portion of these trees shows evidence for higher rates that are consistently shared by a set of genera. Ten genera of the tribe Alsophiini, five genera of Tachymenini, and a clade formed by *Pseudoeryx* and *Hydrops* show rate distributions with a tendency for higher rates of diversification, although median values are not significantly distinct from background rates (Figure S3).

We performed posterior predictive simulations and produced a null distribution of likelihood ratio tests (LRTs) to evaluate whether the BiSSE trait-dependent diversification model is adequate to explain the observed data and if preferring the more complex model is justifiable. Since there is no appreciable difference in parameter estimates and DIC results across the categorizations of color types, we performed simulations based only in the *contrasting* and *cryptic* categories. The posterior predictive simulations using both BiSSE models (trait-dependent and trait-independent) produced trees much smaller than the 594 species of the empirical tree (see left column of Figure 4). Since the stopping criteria for simulations were tree depth, the number of species in each simulated tree was free to vary. Trees simulated under the full model had on average 211 species and only 9% of those showed more species than the observed data. Similarly, trees simulated using the trait-independent model had on average 186 species, of which only 5% were larger than the observed data. With respect to trait frequency, both models simulated datasets biased towards higher frequencies of the *cryptic* color type. On average the full model had 67% and the trait-independent model 84% of the simulations showing frequencies of the *cryptic* color type higher than observed in the empirical data (see right column of Figure 4). Therefore, our results show that both models have similar biases and the characteristics of the data not satisfactorily explained by the BiSSE model are independent to whether rates of diversification are associated with color types.

**Figure 4.**
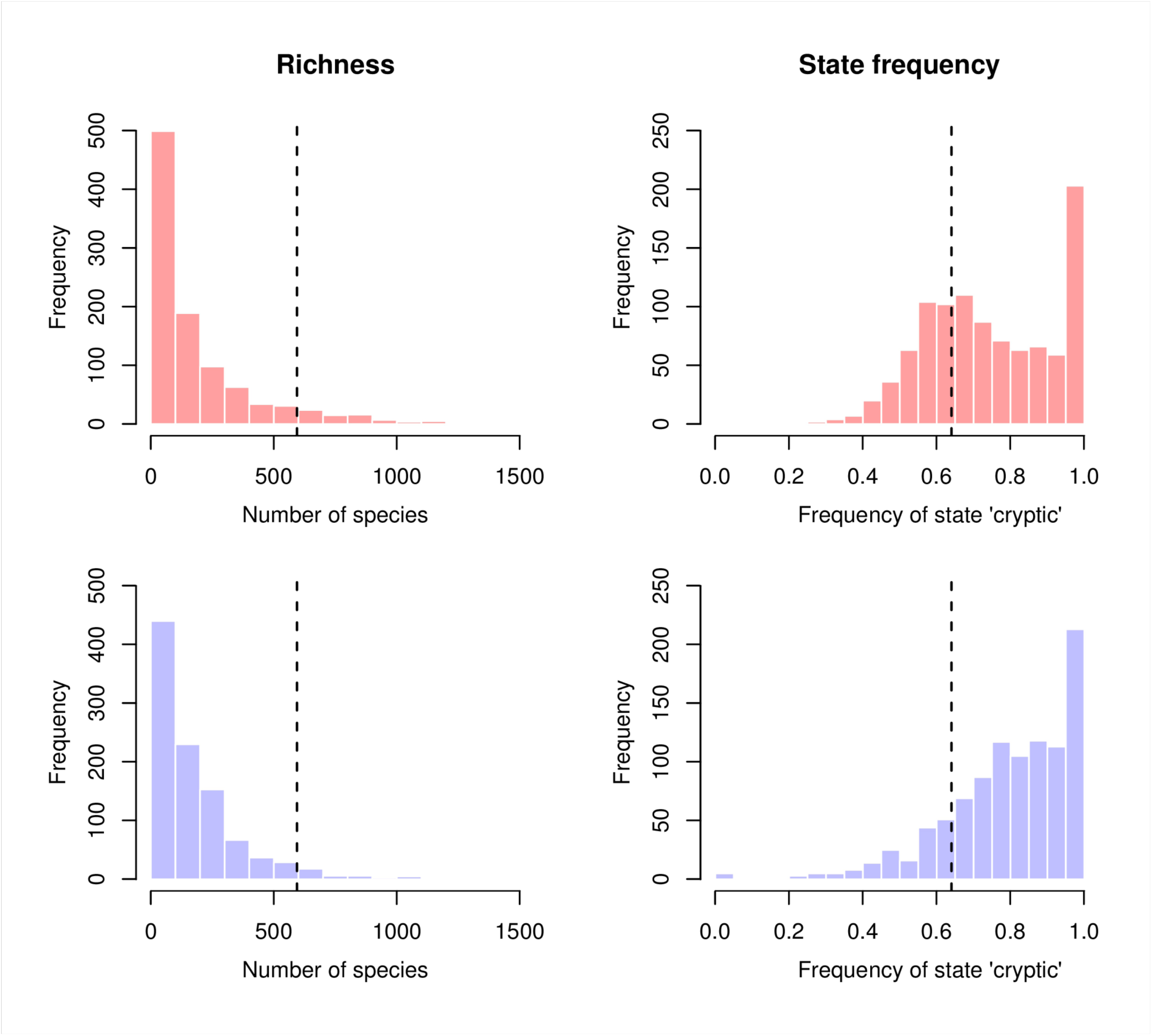
Results from the posterior predictive simulations for BiSSE under the trait-dependent (top row - red) and the trait-independent model (bottom row - blue). Simulation parameters drawn from the joint posterior distribution of each BiSSE model estimated from the observed data (categories *contrasting* and *cryptic*) and phylogenetic trees. At each replicate a phylogeny was simulated under the BiSSE model until the sum of branch lengths from root to tip of the tree was equal to 1. Left column: Total number of species in the resulting phylogenies. Vertical dashed lines show the richness of the Dipsadidae family used in our analysis (594 spp.). Right column: Relative frequency of the state 0 (*cryptic*). Vertical dashed lines show the observed frequency of *cryptic* species in the data (0.64). Note that both BiSSE models show similar deviations from the observed data.

Although parameter estimates under the BiSSE model are robust to our different categorization of color patterns, the ML ancestral state estimate is dependent on whether we consider the *coral-mimic* or *contrasting* color type categories. Ancestral estimates considering species as *coral-mimic* or *non-mimic* suggest the root node had a *non-mimic* color pattern and just two clades show high likelihood for *coral-mimic* ancestors (Figure 1). However, when species are categorized as *contrasting* or *cryptic*, the root node is reconstructed as most likely *contrasting* and several transitions to the *cryptic* form occurred throughout the tree with no evidence of reversals (Figure 1).

## Discussion

### Model selection and (in)adequacy

We fitted the trait-dependent and trait-independent BiSSE models to the data and performed model selection using the Deviance Information Criteria (DIC). DIC showed strong support for the trait-dependent model for both color pattern categorizations (*coral-mimic* versus *non-mimic* and *contrasting* versus *cryptic*). However, our simulations showed that there is no evidence to choose the trait-dependent model over the trait-independent model. Likelihood ratio tests on simulated data consistently gave support to the trait-dependent diversification model (with threshold value of 0.05) despite the fact that data were generated using draws from a binomial distribution, independent of the phylogeny. The BiSSE method is likely to support the trait-dependent model even when there is no real effect of traits on rates of diversification, supporting findings by Rabosky and Goldberg (2015). The choice of the full model by the DIC approach was unjustifiable and might be a statistical artifact. Hence, our preferred model is the trait-independent model in which color type has no influence on diversification.

Posterior predictive simulations show that both the trait-independent and trait-dependent BiSSE models share similar biases. Both BiSSE models are equally inadequate in explaining the observed data whereas the LRT simulations provide evidence that the best model is the simpler model rather than the model in which rates of diversification depend on color type. Results from the Medusa analyses also corroborate this conclusion. There is no support for shifts in rates of diversification and the majority of lineages showing a tendency for higher rates are comprised of cryptic species (Figure S3). At the moment, unlike other models of trait evolution (Pennell et al. 2014), it is not clear which set of summary statistics can be used to assess the adequacy of BiSSE models. More studies are needed to better understand the scenarios under which this and other models of the xxSSE family are prone to misbehave and elect a set of informative summary statistics for predictive posterior checks.

### Uncertainty in ancestral state estimates

The observed variation in ancestral estimates (Figure 1) despite similar parameter estimates for the BiSSE model under both color categorizations (Figures 2 and 3) is most likely an effect of the proportion of states in each of the genera for each categorization. Given our color category definitions, every *coral-mimic* was also included in the *contrasting* category, but the contrary is not true. The proportion of *contrasting* species is higher than *coral-mimics* in several genera and may have increased the likelihood of nodes estimated as *contrasting* when comparing to the same nodes under the *coral-mimic* and *non-mimic* categorization. In both cases changes in color pattern are mostly concentrated within genera, with exception of the tribes Alsophiini and Tachymenini, both comprising only *cryptic* (*non-mimic*) species (marked respectively in green and red in Figure 1). In those tribes the transition from *contrasting* to *cryptic* forms may have occurred at the base of the clades with reversals restricted to the genus *Ialtris* (tribe Alsophiini). These results need to be interpreted with caution, since ancestral estimates are often misleading and a fully resolved tree for the group might falsify our reconstructions by showing reversals from *coral-mimic*/*contrasting* to *non-mimic*/*cryptic* color types present at the genus level.

### Color patterns have no effect on the diversification of dipsadid snakes

The function of coral-mimic and contrasting color patterns in snakes and other groups has been intensely debated since the 50s (Dunn 1954, Hecht and Marien 1956). However, only a few studies have investigated the evolution of snake color patterns using an explicit phylogenetic approach (e.g., Pyron and Burbrink 2009) despite recent advances in comparative methods. Our results show no appreciable effect of the color types in diversification rates despite the impressive diversity of color patterns found in the group. Estimates of diversification rates independent of color types also give support to the trait-independent model, since results do not suggest any concurrence of shifts in diversification rates with the presence of coral-mimic or contrasting patterns. Overall, our results suggest that the ecological functions of *coral-mimic* and *non-mimic* color types in dipsadid snakes have no distinguishable effect on the macroevolution of the group.

The hypotheses of mimicry based on color similarity between mimics and models have received strong support from field experiments and the survival advantage of mimicry is also often demonstrated (Smith 1975, Brodie 1993, Brodie and Janzen 1995, Hinman et al. 1997, Pfennig et al. 2001, Buasso et al. 2006). If mimicry can explain the color types of dipsadid lineages that are similar to coral snakes, the protection attributed to the warning signal can have a positive effect on diversification, similar to the effect of aposematism (Mallet and Joron 1999, Przeczek et al. 2008). However, increased survival has no necessary link to the generation of new species and our results show that distinct color types are not associated with appreciable differences in macroevolutionary patterns in dipsadids, independent of their putative ecological function.

### Contrasting patterns as potential pre-adaptation to coral snake mimicry

Mimetic and contrasting patterns share bright colors that might be significantly more conspicuous to visually oriented predators than cryptic patterns. Field experiments using plasticine replicas provide evidence that the stereotyped alternating, bright colored bands (bearing shades of red, yellow, black and/or white) found among New World elapid species are avoided by predators (Brodie 1993, Brodie and Janzen 1995, Hinman et al. 1997, Pfennig et al. 2001, Buasso et al. 2006), even when these patterns are simplified to an extent that only a single ring occurs on the neck or head and the remaining of the body is plain red (Brodie 1993; see also Hinman et al. 1997). However, contrasting patterns are not restricted to such alternating, bright colored bands. In the absence of mimicry, such patterns might serve as disruptive coloration and deceive predators by creating optical illusions or hindering prey recognition. Indeed, it is plausible that coral-like patterns might function both as mimetic and disruptive coloration (Dunn 1954, Titcomb et al. 2014). The disruptive function (or flicker-fusion effect) can prevent or mitigate maladaptation caused by allopatric distribution of mimics and models, since it provides protection even in the presence of naïve predators. The double function of the contrasting pattern as mimetic and illusionary may help explain the impressive diversity of colors and patterns found among dipsadid snakes. One plausible explanation is that contrasting patterns may serve as pre-adaptations to the evolution of coral snake mimicry in certain lineages.

Ancestral node reconstructions of the *contrasting* pattern in dipsadids show that most genera are likely to have some form of *contrasting* colored ancestor (Figure 1). This pattern changes when considering only the subset of contrasting lineages with color patterns similar to coral snakes. In the latter case, reconstructions show that transitions most likely occurred exclusively within genera in 13 out of 24 genera comprising *coral-mimic* lineages. Among the genera estimated to have *coral-mimic* ancestors 9 out of 11 belong to the Pseudoboini, characterized by a single transition to the *coral-mimic* type (Figure 1). Since *coral-mimics* are a subset of *contrasting* lineages and *contrasting* ancestors occur in nodes deeper in the tree than the *coral-mimics* ancestors, it is likely that *contrasting* patterns not resembling true coral snakes predate the supposed mimetic patterns. The disruptive effect of some contrasting patterns, especially those including light and dark bands, might confer protection against predation and can show rudimentary similarities to potential coral snake models. This scenario could lead to a mimicry relationship as predation pressure can produce a directional selective gradient pulling towards color patterns more similar to those of the model lineage. Although our results suggest this relationship, the current knowledge of the Dipsadidae phylogeny does not provide enough resolution to test this hypothesis, since changes from *contrasting* to *cryptic* patterns occurred almost exclusively within genera.

### Ecological opportunity on the West Indies

We estimated rates of diversification using a model that is agnostic to color types (Medusa) and two clades showed a tendency for elevated rates of diversification, the tribe Alsophiini and part of the tribe Tachymenini (Figure S3). Both tribes comprise cryptic species, with exception of the genus *Ialtris*. The Alsophiini is an interesting case. It most likely dispersed from mainland South America over the ocean to the West Indies, where it is endemic (Maglio 1970, Hedges et al. 2009). The West Indies are known by the remarkable adaptive radiation of *Anoli*s lizards (Losos and Ricklefs 2009) and might also have provided ecological opportunities for the radiation of other squamate reptiles. For instance, Burbrink and colleagues (2012) showed that the diversification of the Alsophiini in the West Indies bears no signal of increased rates of diversification. In contrast, our results show a tendency for increased rates of diversification associated with the same group. Absence of such signal in results by Burbrink and colleagues (2012) may be a reflection of including only the Alsophiini in the analysis whereas herein we used phylogenies including the whole family. The increased rate of diversification of the Alsophiini may be, such as hypothesized by Burbrink et al. (2012), a function of the opportunity for ecological diversification and not associated with the concurrent change from contrasting to cryptic color patterns. It is important to note that the monophyly of the tribe has been contested by some studies (Grazziotin et al. 2012, Pyron et al. 2013) and taxonomic rearrangements are likely to occur in the future. It is possible that tests based on a revised taxonomy and a finer phylogenetic resolution may show support for the hypothesis of ecological opportunity.

## Concluding remarks

Herein we compiled from primary sources and made available a database of color patterns for 594 species of dipsadid snakes, the largest compilation of color descriptions for reptiles to date. We found that *coral-mimic* or *contrasting* patterns have no significant effect on rates of diversification when compared to *non-mimic* or *cryptic* color types. This is an intriguing contrast with the fact that aposematic clades are more species-rich than their cryptic sister groups (Przeczek et al. 2008). Speciation or extinction of mimetic lineages are theoretically linked to the relationship with their models. However, this dependence can be loosened if the mimetic trait is associated with a secondary protective function. Both eventual extinction events caused by allopatry with models and speciation as a result of local adaptation to novel models can be ‘buffered’ by the secondary function of the trait. The protection by illusion might be a precursor for both the remarkable convergence of snake lineages to coral-like forms and the maintenance of mimicry despite the supposed likelihood of mimicry breakdowns.

It is naïve to think that a unique set of traits, such as color patterns, can reflect all relevant factors that drive the dynamics of diversification of any group. A more detailed analysis of our questions could be accomplished by overlapping evolutionary patterns of the Dipsadidae with those of New World coral snakes, for example. However, both phylogenetic data and suitable comparative models are not yet available. Further appreciation of transitions between *contrasting* and *cryptic* color patterns within dipsadid genera can shed light on whether disruptive colors can serve as a pre-adaptation to mimicry and help insert new pieces into the coral snake mimicry puzzle. Understanding under which phylogenetic and ecological scenarios mimicry is likely to evolve is a key factor to explain the patterns of phenotypic convergence observed among distantly related lineages across the tree of life.

## Acknowledgments

We would like to thank Denim Jochimsen, Josef Uyeda, Matt Pennell, Rafael Maia, and Andrew Kraemer for extensive discussions about snakes and statistics. Special thanks to Luke Harmon, DSC advisor, for insightful comments on the work and manuscript. This project applied many ideas discussed at the Phylogenetics Reading Group (PuRGe) at the University of Idaho. DSC was supported by a fellowship from Coordenação de Aperfeiçoamento de Pessoal de Nível Superior (CAPES: 1093/12-6). LRVA was supported by Fundação de Amparo a Pesquisa do Estado de São Paulo (FAPESP, 2012/03038-2). Financial support to PP was provided by Conselho Nacional de Desenvolvimento Científico e Tecnológico (CNPq, 482086/2012-2 and 8256995713198058) and Fundação de Amparo a Pesquisa do Estado do Rio de Janeiro (E-26/110.434/2012 and E-26/111.636/2012). FGG, HZ, and MM also thank FAPESP for a grant (2011/50206-9) and CNPq for fellowships.

## SUPPLEMENTARY INFORMATION

**Figure S1:**
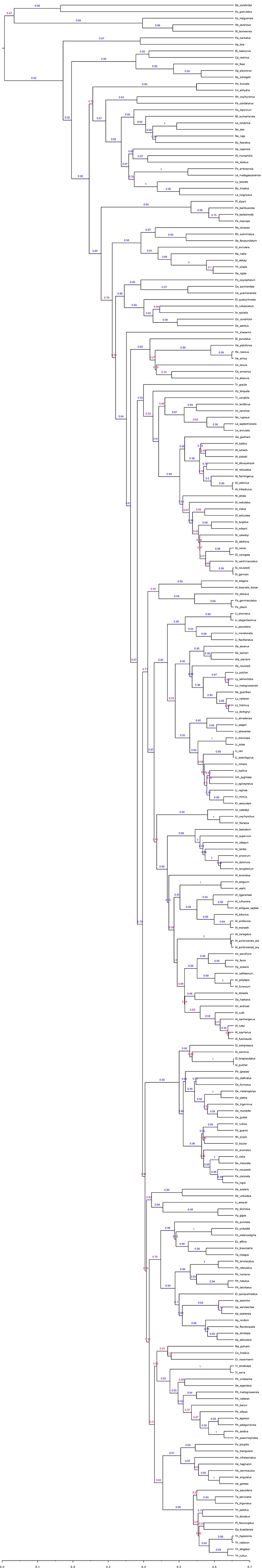
Maximum clade credibility tree with all species used in the analysis. Values in the branches are the respective posterior probabilities for each node. Species are named with the two first letters of the genus followed by the complete species name. Refer to the Table 1 in FigShare (http://dx.doi.org/10.6084/m9.figshare.831493) for the complete name of each species. See Material and Methods of the main text for more information on the methods of tree inference.

**Figure S2:**
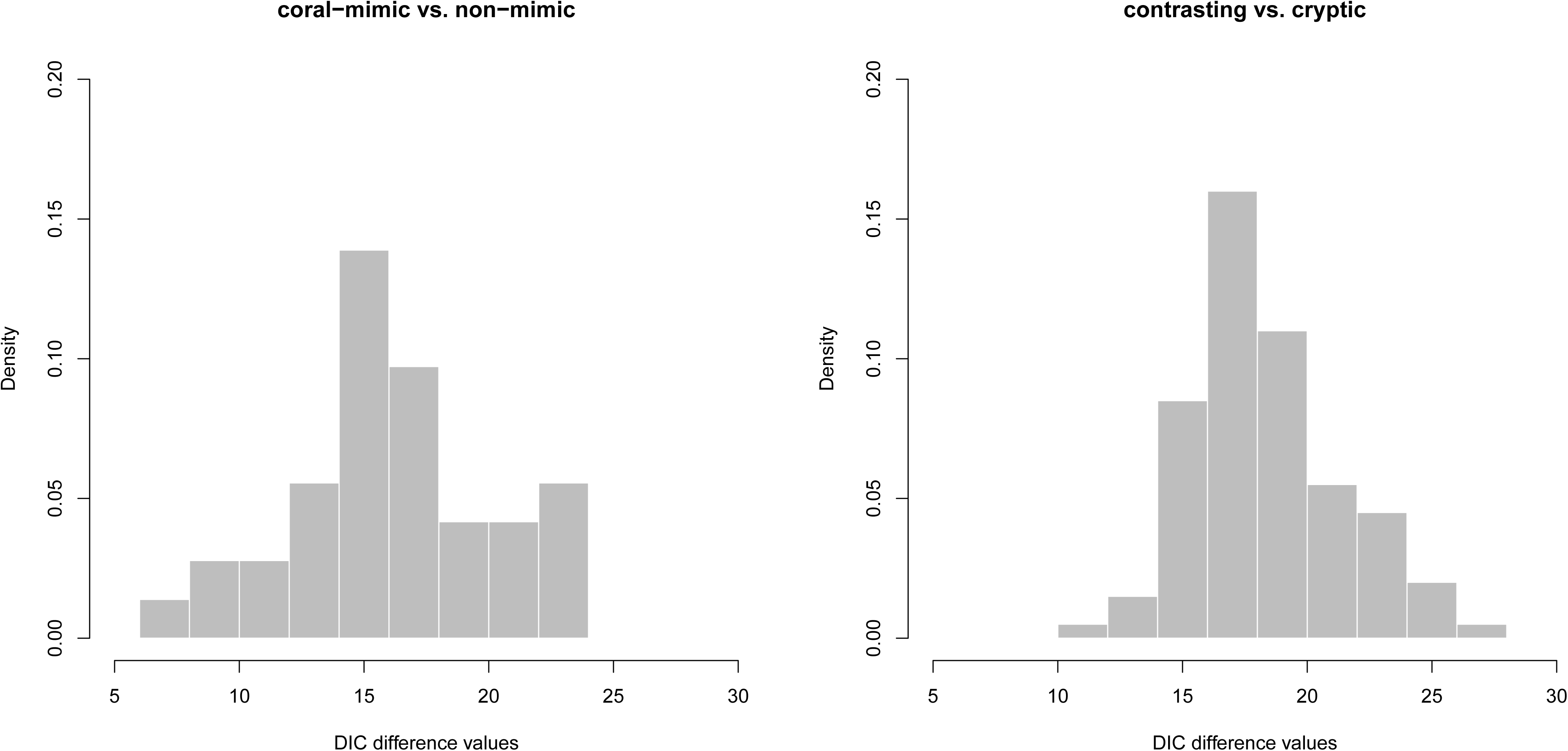
Results of model selection using the Bayesian Deviance Information Criteria (DIC) for the *coral-mimic* versus *non-mimic* (left) and *contrasting* versus *cryptic* (right). DIC values calculated across 100 randomly sampled trees. Values are DIC scores for the trait-dependent model (full model) subtracted from the scores for the trait-independent model (constrained model). Large values—larger than 4 units as a rule of thumb—are expected if the trait-dependent model is to be preferred over the simpler trait-independent model.

**Figure S3:**
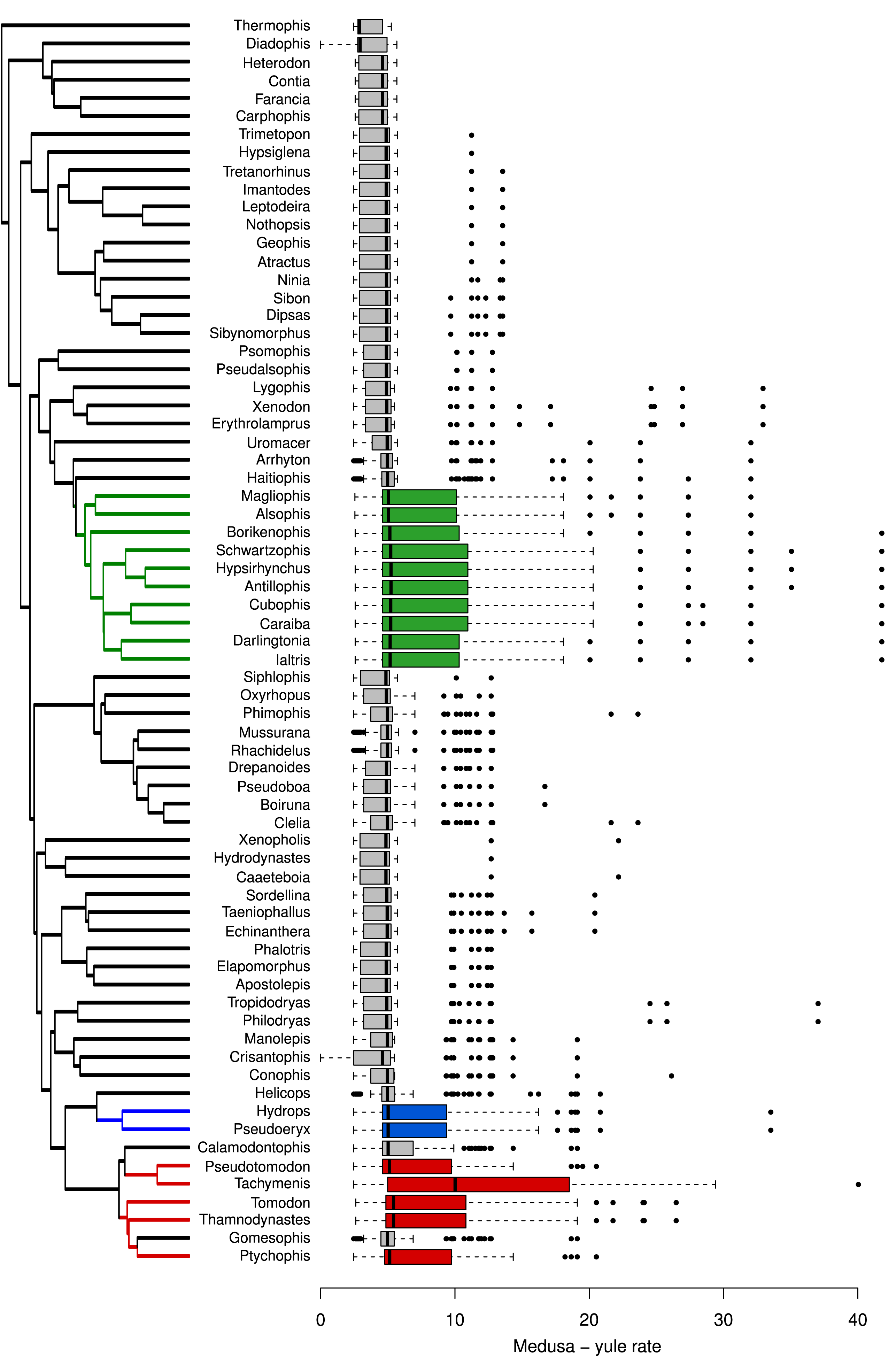
Results of the MEDUSA analyses across 100 trees sampled from the posterior distribution of trees generated by the BEAST analysis. The box plots show the distribution of rates of speciation estimated under a Yule model. The colors green, blue, and red denote genera that show a tendency for higher rates of speciation when compared to the background rates shown in gray. The order of genera and color coding is the same used in the Figure 1. Genera shown in green belong to the tribe Alsophiini, in blue to the tribe Hydropsini, and in red to the tribe Tachymenini.

## References

Alfaro, M. E., F. Santini, C. Brock, H. Alamillo, A. Dornburg, D. L. Rabosky, G. Carnevale, and L. J. Harmon. 2009. Nine exceptional radiations plus high turnover explain species diversity in jawed vertebrates. Proc Natl Acad Sci USA. 106:13410–13414.

Almeida, P. C., D. T. Feitosa, P. Passos, and A. L. C. Prudente. 2014. Morphological variation and taxonomy of *Atractus latifrons* (Günther, 1868) (Serpentes: Dipsadidae). Zootaxa. 3860:64–80.

Benson, D. A., K. Clark, I. Karsch-Mizrachi, D. J. Lipman, J. Ostell, and E. W. Sayers. 2014. GenBank. Nucleic Acids Res. 42:D32–37.

Bittner, T. D., and R. B. King. 2003. Gene flow and melanism in garter snakes revisited: a comparison of molecular markers and island. vs. coalescent models. Biol J Linn Soc. 79:389–399.

Bonnet, X., G. Naulleau, and R. Shine. 1999. The dangers of leaving home: dispersal and mortality in snakes. Biol Conserv. 89:39–50.

Brakefield, P. M. 1990. Genetic drift and patterns of diversity among colour-polymorphic populations of the homopteran *Philaenus spumarius* in a island. archipelago. Biol. J. Linn Soc. 39:219–237.

Brattstrom, B. H. 1955. The coral snake “mimic” problem and protective coloration. Evolution. 9:217–219.

Brodie III, E. D. 1993. Differential avoidance of coral snake banded patterns by free-ranging avian predators in Costa Rica. Evolution. 47:227–235.

Brodie III, E. D., and F. J. Janzen. 1995. Experimental studies of coral snake mimicry: generalized avoidance of ringed snake patterns by free-ranging avian predators. Funct Ecol. 9(2):186–190.

Buasso, C. M., G. C. Leynaud, and F. B. Cruz. 2006. Predation on snakes of Argentina: Effects of coloration and ring pattern on coral and false coral snakes. Stud Neotrop Fauna Environ. 41:183–188.

Burbrink, F. T., S. Ruane, and R. A. Pyron. 2012. When are adaptive radiations replicated in areas? Ecological opportunity and unexceptional diversification in West Indian dipsadine snakes (Colubridae: Alsophiini). J Biogeogr. 39:465–475.

Campbell, J.A. and W.W. Lamar. 2004. The Venomous Reptiles of the Western Hemisphere. 2 Volumes. Cornell University Press, Ithaca, NY.

Downs, F. L. 1967. Intrageneric relationships among colubrid snakes of the genus *Geophis* Wagler. Misc Publ Mus Zool Univ Mich. 131:1–188.

Drummond, A. J., M. A. Suchard, D. Xie, and A. Rambaut. 2012. Bayesian phylogenetics with BEAUti and the BEAST 1.7. Mol Biol Evol. 29(8):1969–1973.

Dunn, E. R. 1954. The coral snake “mimic” problem in Panama. Evolution. 8:97–102.

Endler, J. A. 1986. Natural selection in the wild. Princeton Univ. Press, Princeton, NJ.

FitzJohn, R. G. 2012. Diversitree: Comparative phylogenetic analyses of diversification in R. Methods Ecol Evol. 3:1084–1092.

FitzJohn, R. G., W. P. Maddison, and S. P. Otto. 2009. Estimating trait-dependent speciation and extinction rates from incompletely resolved phylogenies. Syst Biol. 58:595–611.

Forsman, A. 1995. Opposing fitness consequences of colour pattern in male and female snakes. J Evol Biol. 8:53–70.

Gadow, H. 1908. Through southern Mexico. Witherby and Co. Press, London, UK.

Gelman, A., Hwang J., and Vehtari A. 2014. Understanding predictive information criteria for Bayesian models. Stat Comput. 24:997–1016.

Grazziotin, F. G., H. Zaher, R. W. Murphy, G. Scrocchi, M. A. Benavides, Y. P. Zhang, and S. L. Bonatto. 2012. Molecular phylogeny of the New World Dipsadidae (Serpentes: Colubroidea): a reappraisal. Cladistics 28:437–459.

Greene, H. W., and R. W. McDiarmid. 1981. Coral snake mimicry: does it occur? Science 213:1207–1212.

Hecht, M. K., and D. Marien. 1956. The coral snake mimic problem: a reinterpretation. J Morphol 98:335–365.

Hedges, S. B., A. Couloux, and N. Vidal. 2009. Molecular phylogeny, classification, and biogeography of West Indian racer snakes of the Tribe Alsophiini (Squamata, Dipsadidae, Xenodontinae). Zootaxa 2067:1–28.

Hinman, K. E., H. L. Throop, K. L. Adams, A. J. Dake, K. K. McLauchlan, and M. J. McKone. 1997. Predation by free-ranging birds on partial coral snake mimics: the importance of ring width and color. Evolution 51:1011–1014.

Jeffords, M. R., J. G. Sternburg, and G. P. Waldbauer. 1979. Batesian mimicry: field demonstration of the survival value of pipevine swallowtail and monarch color patterns. Evolution 33:275–286.

Katoh, K., K. Kuma, H. Toh, and T. Miyata. 2005. MAFFT version 5: improvement in accuracy of multiple sequence alignment. Nucleic Acids Res. 33:511–518.

King, R. B., and R. Lawson. 1995. Color pattern variation in Lake Erie water snakes: the role of gene flow. Evolution 47:1819–1833.

Lindell, L. E. and A. Forsman. 1996. Sexual dichromatism in snakes: support to flicker-fusion hypothesis. Can J Zool. 74:2254–2256.

Lorioux, S., X. Bonnet, F. Brischoux, and M. De Crignis. 2008. Is melanism adaptive in sea kraits? Amphib-Reptilia. 29:1–5.

Losos, J. B., and R. E. Ricklefs. 2009. Adaptation and diversification on islands. Nature 457:830–836.

Maddison, W. P., P. E. Midford, and S. P. Otto. 2007. Estimating a binary character’s effect on speciation and extinction. Syst Biol. 56:701–710.

Maglio, V. J. 1970. West Indian xenodontine colubrid snakes: their probable origin, phylogeny, and zoogeography. Bull Mus Comp Zool. 141:1–54.

Mallet, J., and M. Joron. 1999. Evolution of diversity in warning color and mimicry: polymorphisms, shifting balance, and speciation. Annu Rev Ecol Syst. 30:201–233.

Mappes, J., N. Marples, and J. A. Endler. 2005. The complex business of survival by aposematism. Trends Ecol Evol. 20(11):598–603.

Marques O. A. V., and Puorto, G. 1991. Padrões cromáticos, distribuição e possível mimetismo em *Erythrolamprus aesculapii* (Serpentes, Colubridae). Mem Inst Butantan. 53(1):127–134.

Martins, M., and M. E. Oliveira. 1993. The snakes of the genus *Atractus* Wagler (Reptilia: Squamata: Colubridae) from the Manaus region, central Amazonia, Brazil Zool Med. 67:21–40.

Martins, M., and M. E. Oliveira. 1998. Natural history of snakes in forests of the Manaus region, central Amazonia, Brazil. Herp Nat Hist. 6:78–150.

Merilaita, S., and J. Lind. 2005. Background-matching and disruptive coloration, and the evolution of cryptic coloration. Proc R Soc Lond B Biol Sci. 272:665–670.

Minin, V., Z. Abdo, P. Joyce, and J. Sullivan. 2003. Performance-based selection of likelihood models for phylogeny estimation. Syst Biol. 52:674–683.

Pennell, M. W., R. G. FitzJohn, W. K. Cornwell, and L. J. Harmon. 2015. Model adequacy and the macroevolution of angiosperm functional traits. Am Nat. 186(2):E33–E50.

Pfennig, D. W., and S. P. Mullen. 2010. Mimics without models: causes and consequences of allopatry in Batesian mimicry complexes. Proc R Soc Lond B Biol Sci. 277:2577–2585.

Pfennig, D. W., W. R. Harcombe, and K. S. Pfennig. 2001. Frequency-dependent Batesian mimicry. Nature 410:323–323.

Pfennig, D. W., C. K. Akcali, and D. W. Kikuchi. 2015. Batesian mimicry promotes pre-and post-mating isolation in a snake mimicry complex. Evolution 69:1085–1090.

Pinheiro, C. E. G. 2011. On the evolution of warning coloration, Batesian and Müllerian mimicry in Neotropical butterflies: the role of jacamars (Galbulidae) and tyrant-flycatchers (Tyrannidae). J Avian Biol. 42:277–281.

Plummer M., N. Best, K. Cowles, and K. Vines. 2006. CODA: convergence diagnosis and output analysis for MCMC. R News. 6:7–11.

Pough, F. H. 1988. Mimicry and related phenomena. In: Biology of Reptilia, vol. 16, Ecology B, Defence and Life History, p. 154–234. Gans, C., Huey, R.B., Eds, New York, Allan R. Liss Inc., New York.

Przeczek, K., C. Mueller, and S. M. Vamosi. 2008. The evolution of aposematism is accompanied by increased diversification. Integr Zool. 3:149–156.

Pyron, R. A., and F. T. Burbrink. 2009. Body size as a primary determinant of ecomorphological diversification and the evolution of mimicry in the lampropeltinine snakes (Serpentes: Colubridae). J Evol Biol. 22:2057–2067.

Pyron, R. A., Burbrink, F. T., and J. J. Wiens. 2013. A phylogeny and updated classification of Squamata, including 4161 species of lizards and snakes. BMC Evol Biol. 13:93.

R Core Team. 2015. R: A language and environment for statistical computing. R Foundation for Statistical Computing, Vienna, Austria. URL http://www.R-project.org/.

Rabosky, D. L., and E. E. Goldberg. 2015. Model inadequacy and mistaken inferences of trait-dependent speciation. Syst Biol. 64:340–355.

Roze, J. A. 1996. Coral Snakes of the Americas: Biology, Identification, and Venoms. Krieger Publishing Company, Malabar, FL.

Santos, J. C., L. A. Coloma, and D. C. Canatella. 2003. Multiple, recurring origins of aposematism and diet specialization in poison frogs. Proc Natl Acad Sci USA. 100:12792–12797.

Savage, J. M. 2002. The amphibian and reptiles of Costa Rica: herpetofauna between two continents, between two seas. Chicago Univ. Press, Chicago, IL.

Savage, J. M., and J. B. Slowinski. 1992. The colouration of the venomous coral snakes (family Elapidae) and their mimics (families Aniliidae and Colubridae). Biol J Linn Soc. 45:235–254.

Sawaya, R. J., O. A. V. Marques, and M. Martins. 2008. Composição e história natural das serpentes de Cerrado de Itirapina, São Paulo, sudeste do Brasil. Biota Neotrop. 8(2):0–0. doi:10.1590/S1676-06032008000200015.

Sazima, I, and A. S. Abe.1991. Habits of five Brazilian snakes with coral-snake pattern, including a summary of defensive tactics. Stud Neotrop Fauna Environ. 26(3):159–164.

Smith, S. M. 1975. Innate recognition of coral snake pattern by a possible avian predator. Science 187:759–760.

Smith, S. A., and C. Dunn. 2008. Phyutility: a phyloinformatics utility for trees, alignments, and molecular data. Bioinformatics 24:715–716.

Speed, M. P., and G. D. Ruxton. 2005. Aposematism: what should our starting point be? Proc R Soc Lond B Biol Sci. 272:431–438.

Speed, M. P., Brockhurst, M. A., and Ruxton G. D. 2010. The dual benefits of aposematism: Predator avoidance and enhanced resource collection. Evolution. 64(6):1622–1633.

Stadler, T. 2009. On incomplete sampling under birth-death models and connections to the sampling-based coalescent. J Theor Biol. 261:58–66.

Stevens, M., and S. Merilaita. 2009. Animal camouflage: current issues and new perspectives. Philos Trans R Soc Lond B Biol Sci. 364:423–427.

Stevens, M., and Ruxton, G. D. 2012. Linking the evolution and form of warning coloration in nature. Proc R Soc Lond B Biol Sci. 279:417–426.

Tanaka, K. 2005. Thermal aspects of melanistic and stripped morphs of the snake *Elaphe quadrivirgata*. Zool Sci. 22:1173–1179.

Thayer, G.H. 1918. Concealing-coloration in the animal kingdom, ed. 2. New York.

Titcomb, G. C., D. W. Kikuchi, and D. W. Pfennig. 2014. More than mimicry? Evaluating scope for flicker-fusion as a defensive strategy in coral snake mimics. Curr Zool. 60:123–130.

Tozetti, A. M., R. B. Oliveira, and G. M. F. Pontes. 2009. Defensive repertoire of *Xenodon dorbignyi* (Serpentes, Dipsadidae). Biota Neotrop. 9:157–163.

Uetz, P., and J. Hosek (eds.). 2014. The Reptile Database, http://www.reptile-database.org/.

Wallace, A. R. 1867. Mimicry and other protective resemblances among animals. Westminster Foreign Q. Rev. 32:1–43.

Wilson, D., R. Heinsohn, and J. A. Endler. 2007. The adaptive significance of ontogenetic colour change in a tropical python. Biol Lett. 3:40–43.

Yule, G. U. 1924. A mathematical theory of evolution, based on the conclusions of Dr J. C. Willis. Philos Trans R Soc Lond B Biol Sci. 213:21–87.

Zaher, H., F. G. Grazziotin, J. E. Cadle, R. W. Murphy, J. C. Moura-Leite, and S. L. Bonatto. 2009. Molecular phylogeny of advanced snakes (Serpentes, Caenophidia) with an emphasis on South American Xenodontines: A revised classification and descriptions of new taxa. Pap Avul de Zool 49:115–153.

Zweifel, R. G. 1960. Herpetology of the Tres Maria Islands. Results of the Puritan-American Museum of Natural History expedition to Western Mexico. Bull Am Mus Nat Hist. 119:77–128.

Zwickl, D. J. 2011. GARLI 2.0 https://www.nescent.org/wg_garli/main_page.

